# A collicular–hypothalamic pathway for social visual awareness

**DOI:** 10.64898/2026.04.17.719268

**Authors:** Kelvin Quinones-Laracuente, Naomi López Caraballo, Asha Y. Caslin, Amy M. LeMessurier, Niles Babin, Robert C. Froemke

## Abstract

Awareness of other individuals is a foundational element of social behavior. Here we examined how specific neural systems detect and signal the visual presence of conspecifics related to social arousal and motivation. We found that visual exposure to videos of other mice can activate hypothalamic oxytocin neurons and promote onset of pup retrieval behavior in naive virgin female mice. A range of social videos depicting conspecifics in diverse contexts, including but not limited to parental behavior, could accelerate onset of pup retrieval compared to non-social controls. Animals would elect to watch social videos over non-social videos. We made photo-tagged recordings from oxytocin neurons of the paraventricular nucleus (PVN), which were preferentially activated during social video viewing. Optogenetic silencing of PVN oxytocin neurons during exposure prevented this behavioral enhancement of pup retrieval onset. We also made photo-tagged recordings from a population of PVN-projecting neurons of the superficial superior colliculus (sSC→PVN units). Compared to other sSC neurons, the sSC→PVN neurons had specialized horizontal direction tuning with robust and sustained responses to social videos. sSC→PVN neurons differentiated visual scenes based on social content, responding most strongly to pup retrieval and less to scenes with increasing numbers of animals. Our results identify a subcortical visual pathway that signals the presence and salience of conspecifics to the oxytocin system, providing a circuit mechanism by which social visual awareness drives neuroendocrine arousal and the acquisition of parental behavior.

## Introduction

The social brain must detect and process the behaviors of other individuals in the environment. Before an animal can interpret social signals, form affiliative bonds, or learn from others, it must first register the presence of relevant conspecifics. This social awareness, i.e., detection and evaluation of other animals, engages arousal systems that modulate attention, motivation, and behavioral state (Adolphs, 2009; Shamay-Tsoory & Abu-Akel, 2016). In many species, visual detection of conspecifics triggers neuroendocrine responses or produces other internal state changes that prepare the organism for various forms of social behavior including parental care and aggression (Insel & Young, 2001; Goodson, 2005; Carcea et al., 2021; Iravedra-Garcia et al., 2025). Yet the neural pathways by which the visual system communicates social presence to subcortical systems governing arousal and social motivation are poorly defined.

The neuropeptide oxytocin, released from the paraventricular nucleus of the hypothalamus (PVN), is a central mediator of social arousal and motivation (Jurek & Neumann, 2018; Froemke & Young, 2021; Grinevich & Neumann, 2021). In mice, oxytocin enables pup retrieval: a parental behavior in which experienced mice respond to the distress calls of isolated pups by carrying them back to the nest (Ehret, 2005). Naive virgin females do not initially retrieve, but acquire the behavior after co-housing with an experienced mother and her litter. Oxytocin released during this experience modulates the left auditory cortex to enhance neural responses to pup distress calls, enabling experienced females to recognize their behavioral significance (Marlin et al., 2015). Importantly, the onset of alloparenting in co-housed virgins is preceded by activation of PVN oxytocin neurons during social interactions with the dam including shepherding and pup retrievals, suggesting that social experience itself triggers the oxytocin signaling that enables behavioral change.

Visual input plays a critical role in this process (Watanabe et al., 2016; Carcea et al., 2021; Ueno et al., 2024; Greer et al., 2025). Virgins observing a retrieving dam through a transparent barrier acquire retrieval behavior, whereas those behind an opaque barrier do not. Retrograde tracing identified a direct projection from the medial superficial layers of the superior colliculus (SC) to PVN oxytocin neurons, and optogenetic activation of this SC→PVN pathway could substitute for visual experience in promoting alloparenting (Carcea et al., 2021). These findings established that a subcortical visual route conveys information about the social environment to the hypothalamic oxytocin system. Whether this pathway signals the mere presence of a conspecific, specific features of social behavior, or the ethological salience of the observed scene remains unknown.

The superior colliculus is an evolutionarily conserved midbrain structure whose superficial layers receive direct retinal input and contain neurons with well-characterized receptive fields and direction selectivity (DePiero et al., 2024; Drager & Hubel, 1976; Wang et al., 2010). The SC has been extensively studied in the context of orienting, prey capture, and defensive behaviors (Basso et al., 2021; Hoy et al., 2019; Shang et al., 2019), but its role in detecting and evaluating social visual stimuli and in routing social information to neuroendocrine circuits that regulate arousal and motivation has not been explored.

Here we developed a head-fixed video playback paradigm to investigate how controlled social visual stimuli engage the SC→PVN→oxytocin pathway and promote parental behavior. We found that visual exposure to conspecifics across a range of social contexts was sufficient to activate hypothalamic oxytocin neurons, drive instrumental seeking of social content, and accelerate the acquisition of pup retrieval. SC neurons projecting to PVN exhibited specialized visual tuning properties and differentiated social scenes based on their content, with responses scaling according to the salience of the social stimulus. These results reveal a subcortical circuit by which visual detection of conspecifics engages the oxytocin system to promote social arousal and enable adaptive parental behavior.

## Methods

All procedures were approved by the New York University Grossman School of Medicine Institutional Animal Care and Use Committee (IACUC).

### Animals

Pup-naive C57BL/6J virgin female mice were bred and raised at NYU Grossman School of Medicine and kept isolated from dams and pups until used for experiments at approximately 8–12 weeks of age. For experiments requiring viral injections, we allowed at least two weeks for viral expression before animals were used. OXT-Cre (Oxt-IRES-Cre) mice on a C57BL/6J background were used for optogenetic tagging and silencing of oxytocin neurons. Naive virgins were pre-screened for retrieval or pup mauling before experiments; around 5% of naive virgins that retrieved at least one pup or mauled pups during pre-screening were excluded from subsequent studies.

### Head-fixation and acclimation

Wild-type C57BL/6J or OXT-Cre virgin females (8-12 weeks at time of surgery) were implanted with a custom L-shaped metal or 3D printed polylactic acid headpost affixed to the skull with dental cement (C&B Metabond). After at least one week of recovery, mice were acclimated to head fixation over two days, beginning with 10–15 min sessions and increasing to 20–30 min. Acclimation was performed in the same behavioral box used for experiments. The head-fixation platform was built using Thorlabs components, consisting of two support towers with extending clamps for headpost fixation and a body tube to restrict movement. Only non-retrieving virgins confirmed by a 10-trial screening were selected for experiments.

### Video stimuli

Videos were recorded using laboratory C57BL/6J mice and pups in a standard behavioral arena at 1080p resolution and 30 fps. Brightness and contrast were adjusted using Adobe Premiere Pro to avoid excessive luminance while achieving sharp, color-contrasted images. Each experimental video consisted of 40 repetitions of a 10-second processed clip, each separated by 5 seconds of dark background, yielding a total presentation of approximately 10 minutes. Videos were displayed on a Feelworld 7-inch monitor (1080p, 1 ms refresh rate, 120 frames per second) positioned 7–8 inches from the mouse’s head at the level of the head-fixation platform.

Luminance was calibrated to a range of 15–20 lux. Six categories of videos were used: 1) Pup retrieval: a dam exiting the nest, walking to an isolated pup, picking it up, and carrying it back to the nest. 2) Isolated conspecific: a virgin female mouse roaming in an empty arena without pups, nest, or other mice. 3) Mother entering nest: a dam approaching and entering the nest without pups present. 4) Retrieval with pup calls: the pup retrieval clip paired with 65–80 kHz ultrasonic pup call playback (40 dB SPL, Tucker-Davis Technologies RZ6) synchronized with the pup being out of the nest. 5) Retrieval error: a dam picking up a pup and placing it outside the nest rather than inside. 6) Dark background: a black screen presented for the same duration as other video conditions.

Two types of abstract control videos were generated from the pup retrieval footage for use in electrophysiology experiments. For PVN recordings, an abstract video was created by computing the average pixel color in each of four quadrants on a frame-by-frame basis, producing a dynamic four-color pattern matched in luminance dynamics. For sSC recordings, a silhouette version was generated by binarizing the retrieval video to produce a black-and-white outline of the moving animals.

### Pup retrieval testing

After each daily video session, virgins were transferred to a free-moving behavioral arena (38 × 30 × 15 cm) prepared with bedding and cotton nesting material and given 20 min to acclimate. Three to five pups (postnatal day 1–5) were then placed in the nest and the mouse was given an additional 5 min of acclimation. For each of 10 trials, a single pup was isolated to the farthest corner from the nest and the mouse was given 2 min to retrieve the pup. If the pup was not retrieved within 2 min, it was manually returned to the nest and the trial was scored as a failure. Retrieval probability for each day was calculated as successful trials divided by total trials. Mice were returned to their home cage after testing. This procedure was repeated once per day for four consecutive days.

### Lever-press instrumental conditioning

To assess active engagement with video stimuli, head-fixed mice were trained to press a lever to self-initiate video presentations. The lever completed a circuit on an Arduino board. Mice were assigned to either a pup retrieval video group (N = 9) or an abstract video group (N = 8). The presses were on a 1:1 schedule for video delivery, and presses during active video playback did not re-trigger the video. Lever presses were recorded daily over seven consecutive sessions. In a separate extinction cohort (N = 6), mice initially pressed for pup retrieval videos for two days, after which the video was switched to the abstract stimulus for days 3–10. Pressing rates (presses per minute) were quantified for each session.

### Surgical procedures

All surgical procedures were performed under isoflurane anesthesia (0.5–2.5%, adjusted based on reflexes and breathing rate) with the mouse placed in a stereotaxic apparatus (Kopf Instruments). Body temperature was maintained at 37 °C throughout surgery with a thermal feedback system. Pre-operative dexamethasone and meloxicam were administered for anti-inflammatory prophylaxis. Post-operatively, buprenorphine extended release (0.05 mg/kg, i.p.) was administered for analgesia. Mice were given at least one week for recovery before any experimental procedures.

### Viral injections

For optogenetic silencing of PVN oxytocin neurons, craniotomies were performed over the PVN (from bregma: −0.72 mm posterior, ±0.12 mm lateral) in OXT-Cre mice. AAV5-FLEX-ArchT was injected at a depth of 5.0 mm using a 5-µl Hamilton syringe with a 33-gauge needle at 0.1 µl/min for a final volume of 1–2.5 µl per side. An optic fiber (400 µm, 0.50 NA, Thorlabs) was implanted 200 µm above PVN on each side and secured with dental cement.

For optogenetic identification of PVN oxytocin neurons, OXT-Cre mice were injected in the PVN with 1 µl of AAV1-CAG::DIO-ChR2 at a titer of 10^13^ vg/ml. After securing the Neuropixels probe, the head was angled to allow insertion of a 200 µm optic fiber to deliver blue light for opto-tagging.

For retrograde identification of sSC→PVN projection neurons, retrograde AAV-ChR2 was injected into the PVN of wild-type C57BL/6J mice. An optic fiber was implanted in the superficial SC to enable photostimulation during Neuropixels recordings. Projection neurons were identified by short-latency responses (<10 ms) to blue light pulses of 5 ms.

### Electrophysiology

Extracellular recordings were performed using Neuropixels 2.0 silicon probes (IMEC) targeted to PVN (from bregma: −0.72 mm posterior, 0.12 mm lateral, 5.0 mm ventral) or sSC (from bregma: −3.08 mm posterior, −0.27 mm lateral, −2.0 mm ventral). The probe was slowly advanced to the target depth and allowed to settle for at least 15 min before recordings began. Recordings were acquired at 30 kHz sampling rate.

### Visual stimulus characterization

To characterize visual response properties of sSC neurons, we presented sparse noise stimuli for receptive field mapping and drifting Gabor patches for direction selectivity measurements. RFs were mapped by presenting brief flashes at a grid of spatial locations across the visual field while recording sSC responses. Direction selectivity was assessed by presenting Gabor patches drifting in eight cardinal directions. The global direction selectivity index (gDSI) was computed for each unit, as described previously (Li et al., 2025; Shi et al., 2017).

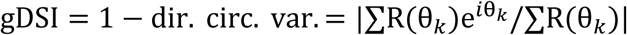

### Opto-tagging

For identification of oxytocin neurons in PVN, blue light pulses (5 ms, 0.5 Hz) were delivered via the implanted optic fiber. Neurons that reliably fired (Z ≥ 5) at short latency (< 10 ms) after the onset of blue light were classified as identified projection neurons. Multiple light intensities were tested to confirm response reliability.

### Spike sorting

Spike sorting was performed using Kilosort 4, followed by manual curation in Phy. Units with refractory period violations > 2% were excluded. Neurons with firing rate less than 0.1 Hz were excluded. For PVN recordings, units were classified as putative oxytocin neurons based on opto-tagging criteria as described above. For sSC recordings, units were classified as sSC→PVN projection neurons based on retrograde ChR2 photostimulation responses.

### Peri-stimulus time histograms

To quantify neural responses to video presentations, spike trains were aligned to the onset of the video stimulus. For PVN recordings, spiking data were binned in 200 ms bins across a window of −2 s to 12 s relative to video onset. For sSC recordings, spiking data were binned in 100 ms bins across a window of −2 to 12 s. Firing rates were converted to z-scores by normalizing each bin to the mean and standard deviation of the pre-stimulus baseline period (all bins preceding 0 s in the respective analysis windows). Population PSTHs were computed by averaging z-scored responses across all units.

### Temporal curve comparisons

To compare neural responses between video conditions across the full stimulus window, we performed point-by-point paired Wilcoxon signed-rank tests at each time bin. P values were corrected for multiple comparisons using the Benjamini-Hochberg false discovery rate (FDR) procedure, with a threshold of q < 0.05. This approach identifies specific temporal windows where population activation significantly diverges between conditions, rather than collapsing responses into a single summary metric.

### Direction tuning analysis

To compare direction selectivity between general sSC and sSC→PVN populations, mean normalized responses to eight cardinal Gabor drift directions were computed for each population. Angular variance across directions was calculated as a measure of tuning selectivity. Pearson correlation was used to assess similarity between the tuning profiles of the two populations.

### Statistical analysis of behavior

For binary retrieval comparisons (percentage of virgins retrieving), Fisher’s exact test was used at each day. For continuous retrieval probability, Wilcoxon rank-sum tests were used for pairwise comparisons between groups at each day. Time to first retrieval across conditions was compared using a log-rank (Mantel-Cox) survival analysis. For lever-press data, a linear mixed-effects model was fitted with group and day as fixed effects and subject as a random intercept (model: Presses ∼ Group × Day + (1|Subject)). Pairwise daily comparisons were performed using Welch’s t-test. All statistical analyses were performed in R and Python (scipy, statsmodels). Significance was set at α = 0.05.

### Histology

At the conclusion of experiments, animals were transcardially perfused with 4% paraformaldehyde. Brains were post-fixed for 1 h at 4 °C, cryoprotected in sequential 15% and 30% sucrose solutions, embedded in OCT compound, and sectioned at 30 µm on a cryostat. Sections were mounted on charged slides and processed for immunohistochemistry as previously described (Valtcheva et al., 2023). Viral expression and probe placement were verified by examining native fluorescence.

## Results

### Video playback of social behaviors accelerates pup retrieval learning

We previously showed that virgin female mice begin retrieving pups faster if given the opportunity to observe experienced mothers through a transparent barrier (Carcea et al., 2021). Here we first asked whether controlled video presentations of social behaviors could similarly promote parental learning. We designed a head-fixed video playback paradigm in which naive virgin mice observed 40 repetitions of 10-second video clips each day for four consecutive days (**Fig. 1A**). After each video session, virgins were tested for pup retrieval in a free-moving arena.

**Figure 1.**
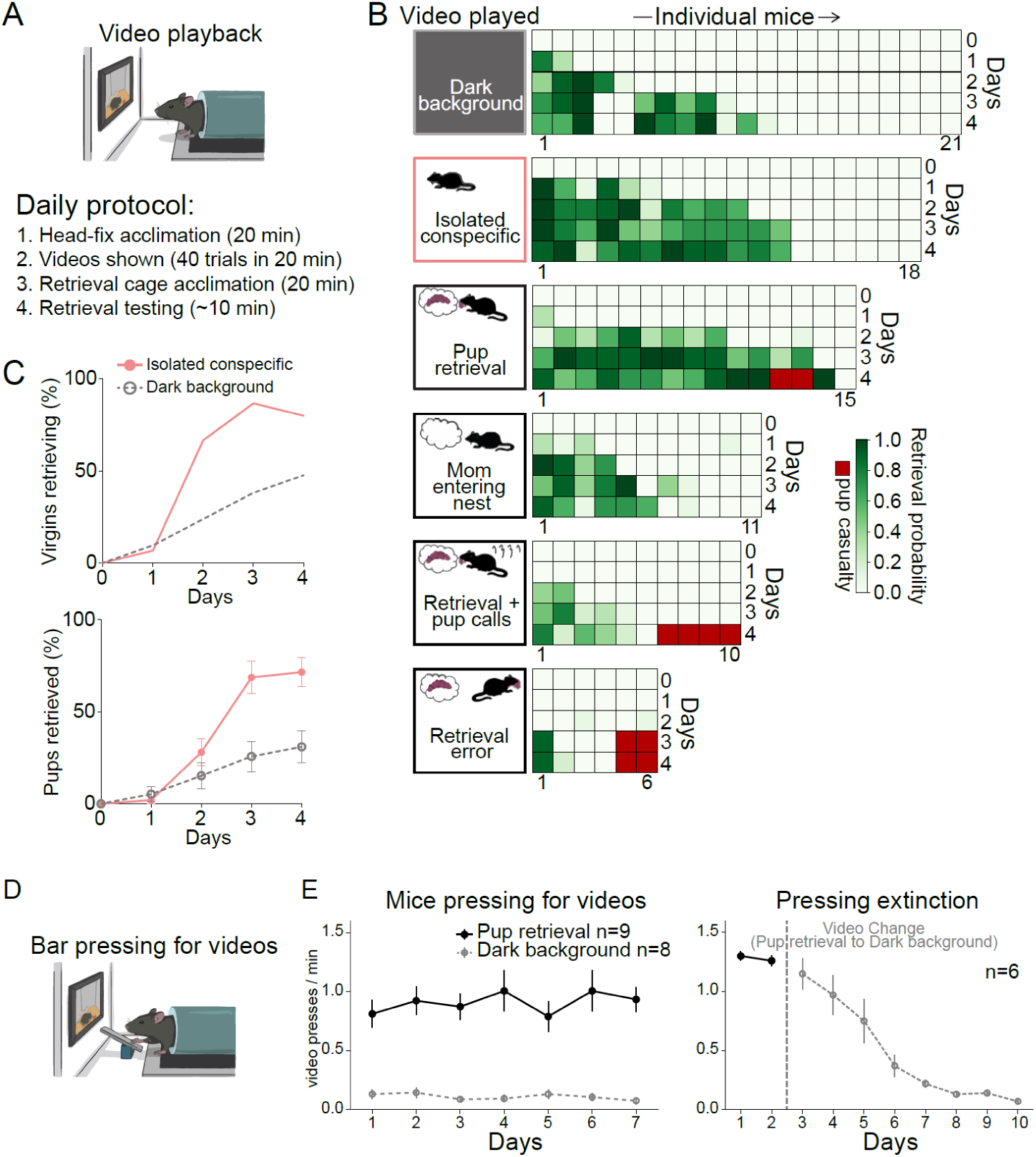
Video playback of social behaviors promotes pup retrieval learning in virgin mice. **(A)** Schematic of the head-fixed video playback and retrieval testing protocol. Each daily session consisted of head-fixation acclimation (20 min), video presentation (40 trials, ∼10 min), free-moving acclimation in the retrieval arena (20 min), and pup retrieval testing (∼10 min). **(B)** Retrieval probability heatmaps for individual virgin mice across 4 days of video exposure. Each column represents one mouse; each row represents one day. Color scale indicates retrieval probability (green) and pup casualties (red). Video conditions shown: dark background (N=21 pup-naïve virgin female mice), isolated conspecific (N=18), pup retrieval (N=15), mother entering nest (N=11), pup retrieval with pup calls (N=10), and retrieval error (N=6). **(C)** Group learning curves showing percentage of virgins retrieving (top) and percentage of pups retrieved (bottom) across days for isolated conspecific (red) and dark background (open circles) conditions. **(D)** Schematic of the lever-press instrumental conditioning paradigm. Head-fixed mice pressed a lever to self-initiate video presentations. **(E)** Left, mean lever presses per day for pup retrieval video (N=9 mice) and abstract video (N=8) conditions. Right, extinction of lever pressing after switching pup retrieval video mice to abstract video presentations. Error bars indicate s.e.m.

We found that video playback of social stimuli accelerated pup retrieval learning compared to dark background controls (**Fig. 1B,C**). Virgins exposed to videos of an isolated conspecific (presented to N=18 mice) began retrieving significantly earlier than dark background controls (N=21; log-rank test, χ² = 7.22, P = 0.0072). By day 2, 66.7% of conspecific-exposed virgins retrieved pups compared to 23.8% of controls (Fisher’s exact test, P=0.017), with the maximal difference on day 3 (86.7% vs. 38.1%, P=0.006). Retrieval probability was also significantly higher in conspecific-exposed virgins by day 2 (Wilcoxon rank-sum, P=0.039) and day 3 (P=0.003). Neither group showed appreciable retrieval on day 1, indicating that a single exposure session was insufficient to induce behavioral change.

We were surprised to find that videos depicting pup retrieval by a dam (N=15) or a mother entering the nest without pups (N=11) promoted retrieval at comparable rates (**Fig. 1B**). In contrast, pup retrieval was less successfully acquired by virgins exposed to videos of retrieval paired with pup call audio (N=10) or videos of incorrect retrieval in which the dam in the movie placed the pup outside the nest (N=6; **Fig. 1B**). These results demonstrate that virtual social experience, even without direct physical interaction, is sufficient to promote parental behavior, and that a range of social stimuli, not exclusively parental demonstrations, can promote this learning.

To assess whether mice actively choose to observe social videos, we trained head-fixed mice to press a lever for video playback (**Fig. 1D**). Mice pressed significantly more for pup retrieval videos (N=9 mice) than for abstract videos (N=8; linear mixed-effects model, group effect: z = 5.20, P < 0.001; **Fig. 1E**). This preference was stable across seven days of testing (all daily pairwise comparisons P < 0.001, Welch’s t-test). When mice previously pressing for retrieval videos were switched to abstract stimuli, pressing rates extinguished over approximately one week, declining from 1.3 to 0.07 presses/min (**Fig. 1E**). These findings indicate that mice find social video content intrinsically reinforcing and can discriminate social from non-social visual stimuli.

### Hypothalamic oxytocin neurons are recruited by social video playback

We previously found that sSC neurons project to oxytocin neurons in mouse PVN, and that optogenetic stimulation of this projection could replace the effect of visual exposure to retrievals when virgins were then subsequently tested on pup retrieval themselves. Here we next asked if PVN neurons respond to virtual social stimuli. We performed Neuropixels 2.0 silicon probe recordings in the PVN of head-fixed virgin mice during video presentations (**Fig. 2A**). Individual PVN neurons showed sustained activation during pup retrieval videos that was absent or diminished during abstract control videos (**Fig. 2B**). At the population level (n=92 single-units), pup retrieval videos evoked significantly greater activation than abstract stimuli, with prominent divergence emerging during two temporal windows: an early-mid epoch (5.1–5.7 s post-onset) and a late sustained epoch (7.1–8.9 s; point-by-point paired Wilcoxon signed-rank tests with Benjamini-Hochberg FDR correction, q < 0.05; **Fig. 2C**). These temporally precise windows of divergence indicate that PVN population dynamics are modulated by specific phases of the unfolding social behavior depicted in the video, rather than exhibiting a uniform increase in activity.

**Figure 2.**
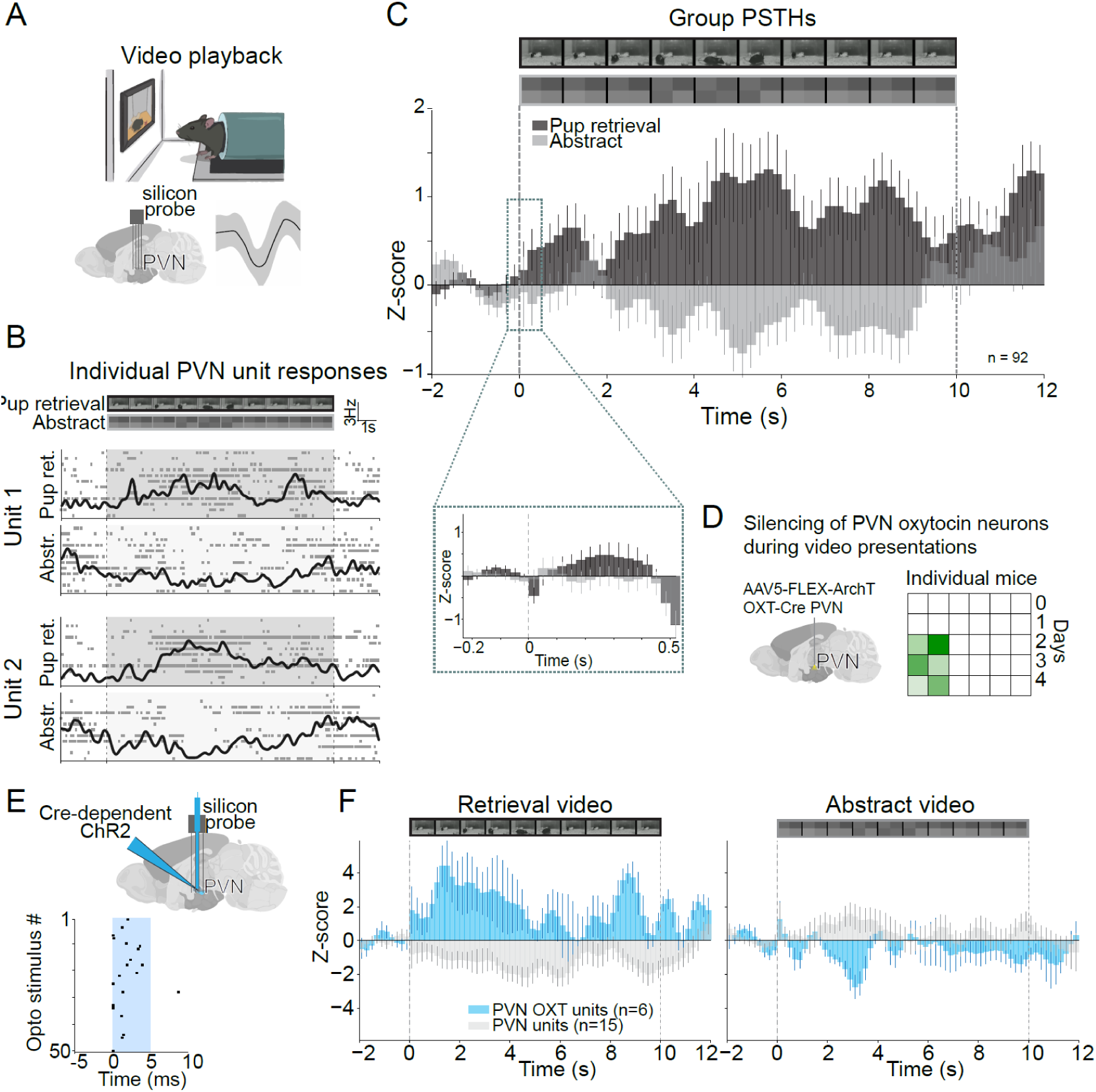
PVN oxytocin neurons respond to social videos and enable for retrieval learning. **(A)** Schematic of silicon probe recordings in the paraventricular nucleus of the hypothalamus (PVN) during head-fixed video playback. **(B)** Example single-unit responses from two PVN neurons to pup retrieval (black) and abstract (gray) video presentations. background, raster plots; foreground, peri-stimulus time histograms (PSTHs). **(C)** Group PSTH of PVN neurons (n=92 single-units) during pup retrieval (black) and abstract (gray) video playback. Video frames shown above. Shading indicates s.e.m. PVN neurons showed sustained activation throughout the pup retrieval video relative to abstract stimuli. **(D)** Optogenetic silencing of PVN oxytocin neurons during pup retrieval video presentations. AAV5-FLEX-ArchT was injected into the PVN of OXT-Cre mice. Green light was delivered concurrently with video playback. Heatmap shows retrieval probability for individual animals (N=6 mice) across 4 days of exposure to pup retrieval videos with concurrent oxytocin neuron inhibition. **(E)** Opto-tagging of PVN oxytocin neurons. Cre-dependent ChR2 was injected into the PVN of OXT-Cre mice and neurons were identified by short-latency responses (<10 ms) to blue light photostimulation of 5 ms during silicon probe recordings. Bottom, example photostimulation response. **(F)** PSTHs of opto-tagged PVN oxytocin neurons (cyan, n=6 units) and other PVN units (gray, n=15 units) during pup retrieval (left) and abstract (right) video playback. Video frames shown above each PSTH.

We next asked whether oxytocin neuron activity during video exposure is required for video-induced retrieval learning. We injected AAV5-FLEX-ArchT into the PVN of OXT-Cre mice and delivered green light bilaterally during video presentations to silence oxytocin neurons (**Fig. 2D**). Optogenetic suppression of PVN oxytocin neurons during pup retrieval video exposure prevented the acquisition of retrieval behavior (N=6 mice; **Fig. 2D**), similar to the requirement of oxytocin signaling for live visual exposure to enhance virgin pup retrieval onset. Thus, PVN oxytocin neuron activity during virtual social experience is necessary for this form of parental learning.

To ask how identified oxytocin neurons contribute to the video-evoked PVN responses, we used Cre-dependent ChR2 in OXT-Cre mice to opto-tag oxytocin neurons during silicon probe recordings (**Fig. 2E**). Identified PVN oxytocin neurons (n=6 units) showed preferential activation during pup retrieval videos compared to abstract stimuli, while other simultaneously recorded PVN units (n=15) had weaker selectivity (**Fig. 2F**). These results establish that hypothalamic oxytocin neurons are selectively recruited by virtual social stimuli and that their activity is required for video-induced parental learning.

### Visual features encoded by sSC→PVN projections

Retrograde tracing identified monosynaptic projections from sSC to PVN oxytocin neurons (Carcea et al., 2021). We characterized visual response properties and video-evoked activity of sSC neurons, including those projecting to PVN, using Neuropixels 2.0 recordings in sSC (**Fig. 3A**). SC neurons responded robustly to light stimulation (**Fig. 3B**) and exhibited well-defined receptive fields (**Fig. 3C,E**; n=1,286 units). Population-level direction tuning to drifting Gabor stimuli was broadly distributed across orientations (**Fig. 3D,F**; n=1,690 units, gDSI ≥ 0.2).

**Figure 3.**
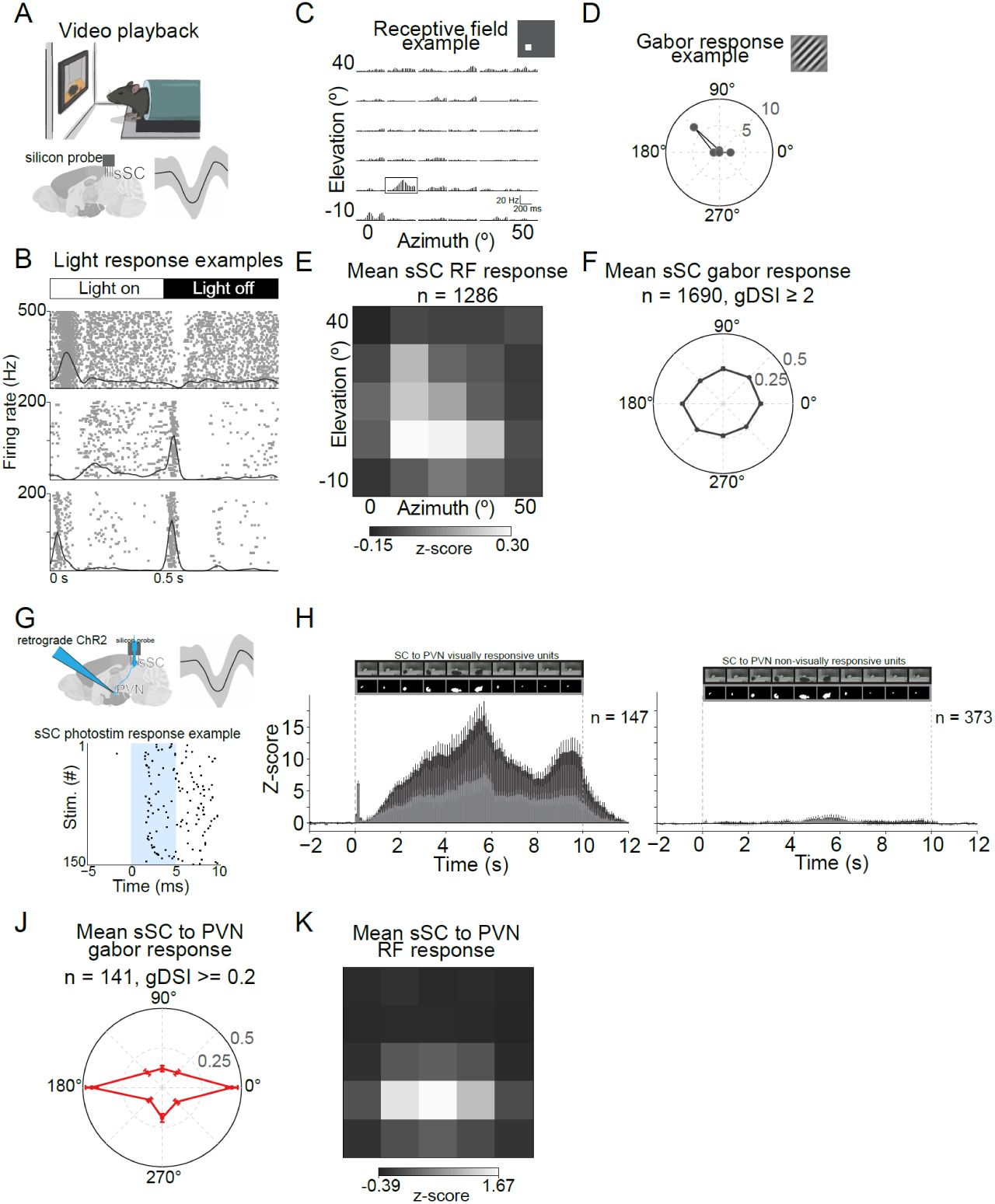
sSC→PVN neurons are tuned to social visual features of conspecific motion. **(A)** Schematic of silicon probe recordings in sSC during head-fixed video playback. **(B)** Example light on/off responses of sSC neurons, demonstrating visual responsiveness. background, raster plot; foreground, PSTH. **(C)** Example receptive field (RF) map of an sSC neuron obtained using sparse noise stimulation. Color scale indicates z-scored firing rate across azimuth and elevation. **(D)** Example direction selectivity of an sSC neuron measured with drifting Gabor stimuli. Polar plot shows normalized response magnitude as a function of stimulus direction. **(E)** Mean RF response across the sSC population (n=1,286 units). **(F)** Mean direction tuning of sSC neurons (n=1,690; global direction selectivity index [gDSI] ≥ 0.2), shown as a polar plot. **(G)** Identification of sSC neurons projecting to PVN. Retrograde AAV-ChR2 was injected into the PVN and silicon probe recordings were performed in the sSC with photostimulation. Bottom, example photostimulation response showing short-latency activation of an sSC→PVN projection neuron. **(H)** Group PSTHs of sSC→PVN projection neurons during video playback, separated by visual responsiveness. Left, visually responsive sSC→PVN neurons (n=147) showed robust activation during video presentations. Right, non-visually responsive sSC→PVN neurons (n=373) showed minimal modulation. Video frames shown above. **(J)** Mean direction tuning of sSC→PVN projection neurons (n = 141; gDSI ≥ 0.2), shown as a polar plot. **(K)** Mean RF response of sSC→PVN projection neurons. Color scale indicates z-scored firing rate.

We identified sSC→PVN projection neurons by injecting retrograde AAV-ChR2 into the PVN and performing photostimulation during sSC recordings (**Fig. 3G**). Visually responsive sSC→PVN neurons (n=147 units responding to light stimulation) also showed strong activation during video playback, whereas non-visually responsive sSC→PVN neurons (n=373) were unresponsive to videos (**Fig. 3H**). Comparing the full time-course of responses, sSC→PVN neurons were much more strongly activated during pup retrieval videos than abstract silhouette controls, with significant divergence beginning ∼1.25 s post-stimulus onset and persisting throughout the remainder of the stimulus (99 of 120 bins, FDR q < 0.05; **Fig. 3H**).

Notably, sSC→PVN projection neurons exhibited a distinct tuning architecture compared to the general sSC population. While the broad sSC population responded nearly uniformly to all Gabor drift directions (angular variance: 0.0004, r=0.65, P=0.08 tuning selectivity), the sSC→PVN subpopulation showed pronounced selectivity for horizontal motion axes (0° and 180°; angular variance: 0.018, a 45-fold increase; **Fig. 3F,J**). These data show that the SC→PVN pathway filters visual space to extract specific relevant motion features of conspecific (i.e., social) motion.

### sSC→PVN neurons differentially encode social visual content

Finally, we asked how sSC→PVN neurons might distinguish between social scenes of different content. We compared population responses (n=192 sSC→PVN projection units) to four video types: pup retrieval, a single mouse, two mice together, and three mice together (**Fig. 4A–D**). sSC→PVN neurons showed the strongest responses to pup retrieval videos, with lower activation as the number of animals in the scene increased. Using point-by-point comparisons against the pup retrieval baseline (paired Wilcoxon signed-rank tests, FDR-corrected), we found that all three alternative conditions diverged significantly from pup retrieval in activation magnitude. The single mouse video separated within 50 ms of onset (70 of 120 bins, q < 0.05), while two-mouse and three-mouse videos diverged at 350 ms and 950 ms respectively, with widespread separation across 89% and 85% of the stimulus window (**Fig. 4A–D**). These results reveal a specificity for the subset of sSC neurons projecting to the hypothalamus, in which sSC→PVN neurons differentiate social from non-social stimuli and also have graded responses based on social content of the visual scene.

**Figure 4.**
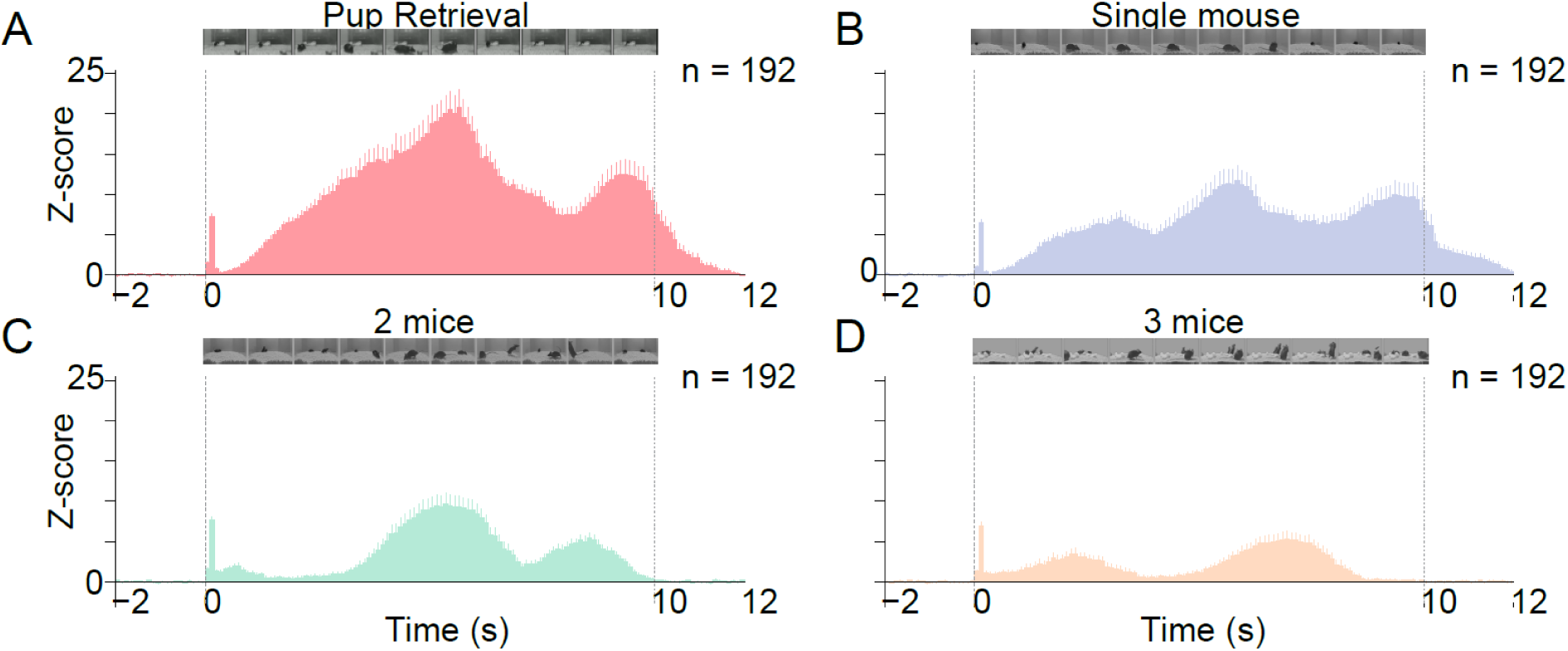
sSC→PVN projection neurons respond more strongly to single adults. (A–D) Group PSTHs of sSC→PVN projection neurons (n=192 units) in response to videos depicting different social scenes. **(A)** Pup retrieval by the mother. **(B)** A single mouse in the arena. **(C)** Two mice interacting. **(D)** Three mice in the arena. Film strip frames from each video shown above the corresponding PSTH. sSC→PVN neurons showed the strongest responses to pup retrieval and single mouse videos, with diminishing responses as the number of animals in the scene increased. Shading indicates s.e.m.

## Discussion

We showed that head-fixed video playback of social behaviors promoted pup retrieval learning in naive virgin mice. This emulated and extends our past work on accelerating alloparenting onset through co-housing or exposure to experienced animals performing maternal caregiving behaviors (Marlin et al., 2015; Schiavo et al., 2020; Carcea et al., 2021). Replacing live exposure with video presentation is experimentally advantageous. By decoupling the visual stimulus from the physical presence of a demonstrator, it enables precise control over the content, timing, and repetition of social experience while eliminating confounding olfactory, auditory, and tactile cues. As mice actively sought social video content through instrumental lever pressing, this suggests that these stimuli are salient and/or rewarding leading to increased viewing compared to the lower number of lever presses and extinction of lever press behavior when animals viewed abstracted videos.

As a range of social videos (not exclusively parental retrieval) accelerated learning, this suggests that the relevant visual signal is the presence and movement of a conspecific rather than a specific behavioral sequence. We hypothesize this leads to a heightened arousal state in the observer, enhancing her performance and learning of pup retrieval when tested shortly after exposure (as opposed to observational learning and copying behavior from the videos). This indicates that the PVN oxytocin system is sensitive to specific patterns of visual input tracking the statistics of conspecific visual features and motion, rather than by recognition of a specific behavioral template. However, the failure of retrieval error videos to promote learning suggests that not all social stimuli are equivalent, and that the outcome of the observed behavior might also be important. In these cases, a minority of animals did seem to copy the observed behavior instead, opening the possibility for bona fide observational learning and behavioral mimicry in mice in some cases. Virtual social experience engaged a subcortical visual pathway from sSC to PVN oxytocin neurons These sSC→PVN projection neurons exhibited specialized direction tuning and had graded responses according to the social content of the visual scene, with the strongest activation evoked by single parents engaged in pup retrieval. The sSC→PVN pathway we characterized provides a plausible substrate for rapid, pre-attentive routing of social visual information to the neuroendocrine system. The horizontal motion selectivity of sSC→PVN neurons corresponds to approach and retreat which are behavioral vectors with high ethological relevance for rodent social interactions (Shang et al., 2019; Solié et al., 2022). The progressive decrease in sSC→PVN activation from pup retrieval to single-mouse to multi-animal scenes suggests that these neurons may be tuned to the salience or identifiability of individual social agents, rather than to aggregate visual motion. This is consistent with proposals that the sSC prioritizes detection of discrete, behaviorally relevant objects in the visual field (Basso et al., 2021).

Social learning is disrupted in neurodevelopmental and psychiatric conditions including autism spectrum disorder and schizophrenia spectrum and other psychotic disorders, which are also associated with atypical oxytocin signaling and altered visual attention to social stimuli (Chevallier et al., 2012; Meyer-Lindenberg et al., 2011). The sSC→PVN pathway described here may represent a subcortical conserved circuit for social cognition whose dysfunction could contribute to these conditions. While we focused on the sSC→PVN projection, other visual pathways including geniculocortical circuits likely contribute to social visual processing (Goltstein et al., 2021; Pessoa & Adolphs, 2010). Our results demonstrate that virtual social experience can engage the same neuroendocrine circuits as live social interaction, opening new avenues for studying the mechanisms of social learning with the precision afforded by new computational tools.

## Acknowledgements

This work was supported by grants from the NIH (NYU-H+H CTSI grant UL1TR001445) and the Leon Levy Scholarships in Neuroscience to K.Q.-L.; an NSF Graduate Research Fellowship and HHMI Gilliam Fellowship to N.L.C.; an NYU CTSA T1 Predoctoral Fellowship and NIH grant NS139546 to A.C.; NIH grants MH019524, MH123016, and DC021581 to A.M.M.; and NIH grants HD088411, NS107616, and NS138066 to R.C.F.

